# The airborne transmission of viruses causes tight transmission bottlenecks

**DOI:** 10.1101/2023.04.14.536864

**Authors:** Patrick Sinclair, Lei Zhao, Clive Beggs, Christopher J. R. Illingworth

**Affiliations:** MRC University of Glasgow Centre for Virus Research, Glasgow, UK; Section for GeoGenetics, Globe Institute, University of Copenhagen, Copenhagen, Denmark; Carnegie School of Sport, Leeds Beckett University, UK

## Abstract

The transmission bottleneck describes the number of viral particles that found an infection in a new host. Previous studies have used genome sequence data to suggest that transmission bottlenecks for influenza and SARS-CoV-2 involve few viral particles, but the general principles underlying these bottlenecks are not fully understood. Here we show that, across a broad range of circumstances, tight transmission bottlenecks arise as a consequence of the physical process underlying airborne viral transmission. We use a mathematical model to describe the process of infectious particles being emitted by an infected individual and inhaled by others nearby. The extent to which exposure to particles translates into infection is determined by an effective viral load, which is calculated as a function of the epidemiological parameter R_0_. Across multiple scenarios, including those present at a superspreading event, our model suggests that the great majority of transmission bottlenecks involve few viral particles, with a high proportion of infections being caused by a single viral particle. Our results provide a physical explanation for previous inferences of bottleneck size and predict that tight transmission bottlenecks prevail more generally in respiratory virus transmission.

## Introduction

The transmission bottleneck describes the number of viruses establishing an infection in a newly infected host. The transmission bottleneck has an important influence upon virus evolution: The tighter the bottleneck, the less genetic diversity will be transmitted between individuals. Genomic data has shown that for influenza and SARS-CoV-2 infection, the transmission bottleneck generally involves few viral particles^1–4^. This tight bottleneck limits the role of within-host evolution; variants need to be generated *de novo* before selection at that scale can take effect^5^.

A variety of experimental and statistical approaches have been used to infer bottleneck sizes from genomic data. In animal models, barcoded viruses allow for a straightforward identification of the number of viruses founding a population^6^. Where barcoding is not possible, deep sequencing of a viral population can be used to assess the appearance or non-appearance of minor variants following the bottleneck^7^, or changes in variant frequencies during the transmission process^8–11^. The statistical estimation of bottlenecks requires care, with falsely called variants having the potential to increase the bottleneck size inferred^12,13^: errors in processing data have led to widely disparate claims of bottleneck size. The size of the transmission bottleneck arises from the physical process underlying viral transmission^14,15^. To form part of the bottleneck, a virus must travel from one host to another, then infect a host cell so that its descendants spread and initiate an infection. Here we examine the hypothesis that the specific physical processes underlying airborne virus transmission favour a tight population bottleneck. Rather than considering a particular genomic dataset, we aim to outline a general solution to the problem of bottleneck inference in respiratory virus transmission.

Studies of the physical process of disease transmission have a long history^16^. Although debates around droplet-based versus aerosol-based transmission^17^ have caused some controversy^18^, some things are clear. Coughing, speaking and sneezing each emit a broad distribution of particle sizes^19–21^. Emitted particles are affected by evaporation, sedimentation and diffusion^22^. Ventilation removes particles from the air, while in the absence of immediately finding a new host, viruses in emitted particles begin to decay^23^. The SARS-CoV-2 pandemic inspired a body of work assessing the risk of infection in a broad range of scenarios^24–27^. We here build upon this literature to assess the expected transmission bottleneck for infections under a variety of scenarios. Our results provide a strong expectation, independent of genome sequence data, that most events of respiratory virus transmission will involve a tight population bottleneck.

## Results

A physical model suggested that where viruses spread via airborne transmission, the resulting transmission bottlenecks are likely to be small. In our model, infectious particles with a distribution of sizes are emitted by an infected individual in regular coughing events. Particles spread through the environment by diffusion. They are subject to evaporation and loss from the air by ventilation and sedimentation. Viruses in emitted particles have a half- life, and degrade over time (Figure 1A). Infection occurs when an uninfected individual is exposed to infectious particles. Each individual inhales a stochastic number of particles; the precise number of particles of each size is modelled as a random emission from a distribution based on a mean level of exposure (Figure 1B). The number of viruses founding an infection is then calculated as the result of a second stochastic process. The particles inhaled represent a cumulative volume of liquid, considered in terms of the volume at the point of emission. Particles have an effective viral load, describing the mean number of viruses per unit volume that would be expected to found infection. The transmission bottleneck is then modelled as a random variable with mean equal to the product of these two values (Figure 1C). Transmission was simulated in environments with different prevailing conditions, describing an office, a nightclub, a bus, and a domestic living room. Distributions of bottleneck sizes were calculated across repeated simulations of each environment.

**Figure 1:**
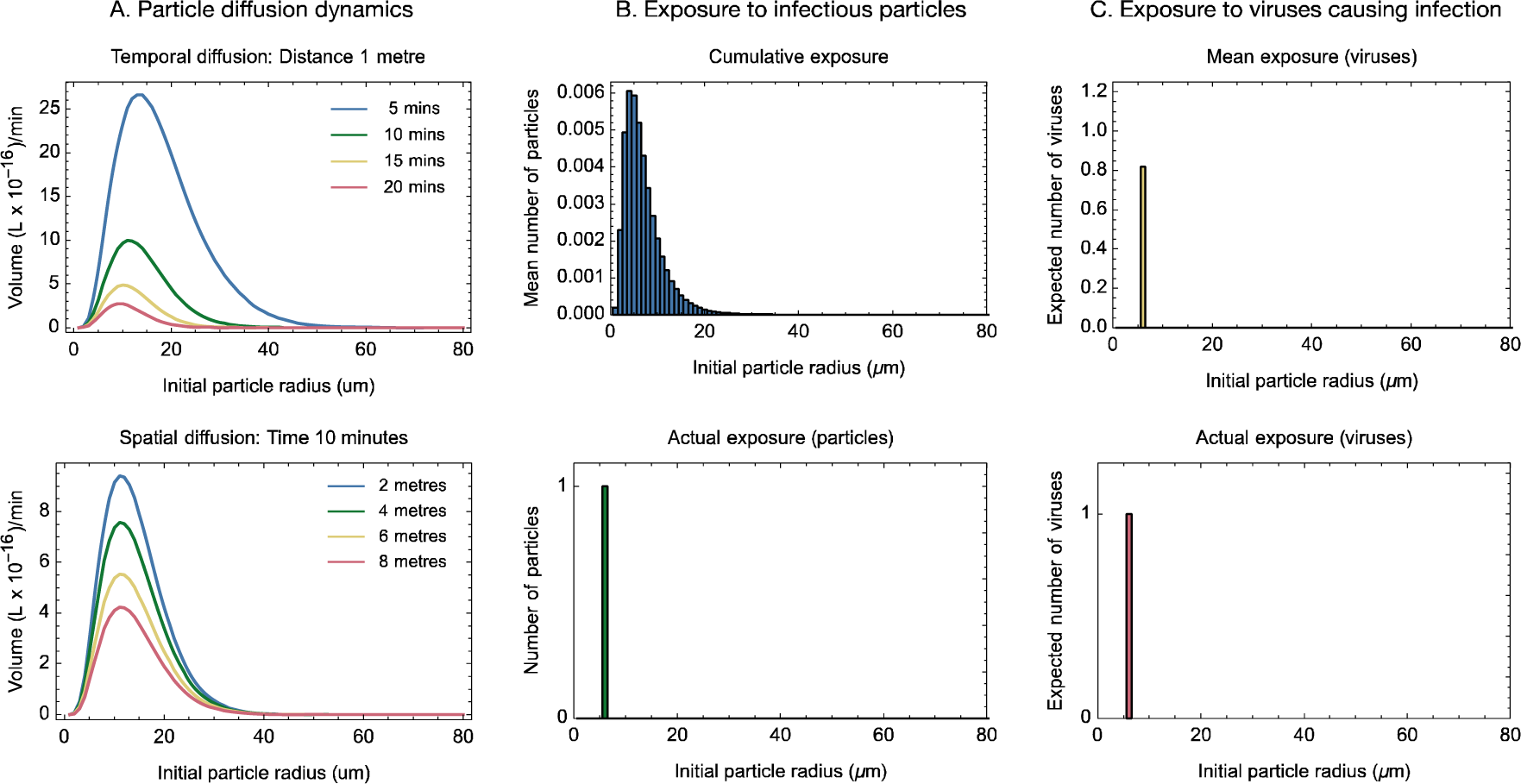
Method for simulating transmission events. **A.** A computational model described the emission and subsequent dynamics of virus-containing particles following a single cough. We modelled the diffusion of particles of different sizes through space and time, accounting for evaporation, sedimentation, ventilation, and the degradation of viruses within infectious particles. Our model describes an effective volume of material, quantifying the level of exposure at a given time and location in terms of an equivalent volume of material at the time of emission, prior to evaporation or virus degradation. Data shown describe particle dynamics in an office environment. **B.** For any given individual in an environment, the cumulative exposure describes the expected number of particles of each size inhaled by an individual during the time period of the model. The actual number of particles inhaled by an individual was then modelled as a set of Poisson random samples parameterised by the cumulative exposure. In this example a person was exposed to one particle of initial radius 6μm. Particle size is here measured at the time of emission, prior to evaporation. **C.** The mean number of viruses founding infection was described as a function of the particles inhaled, and an effective viral load, which describes the expected number of viruses per ml of liquid, that will go on to found infection. Here the expected number of viruses is approximately 0.8. The actual number of viruses founding infection was calculated as a set of Poisson random samples parameterised by the mean number of viruses founding infection. Infection occurred if at least one virus founded an infection. In this case the transmission bottleneck involved a single virus.

Running our model under default parameters suggested that tight transmission bottlenecks would predominate in each of our environments, with roughly 80% of transmission events involving ten or fewer viruses (Figure 2). Distributions often had a long tail: Across 10^5^ simulations of the office environment we identified three cases involving a transmission bottleneck of more than 200 viruses.

**Figure 2:**
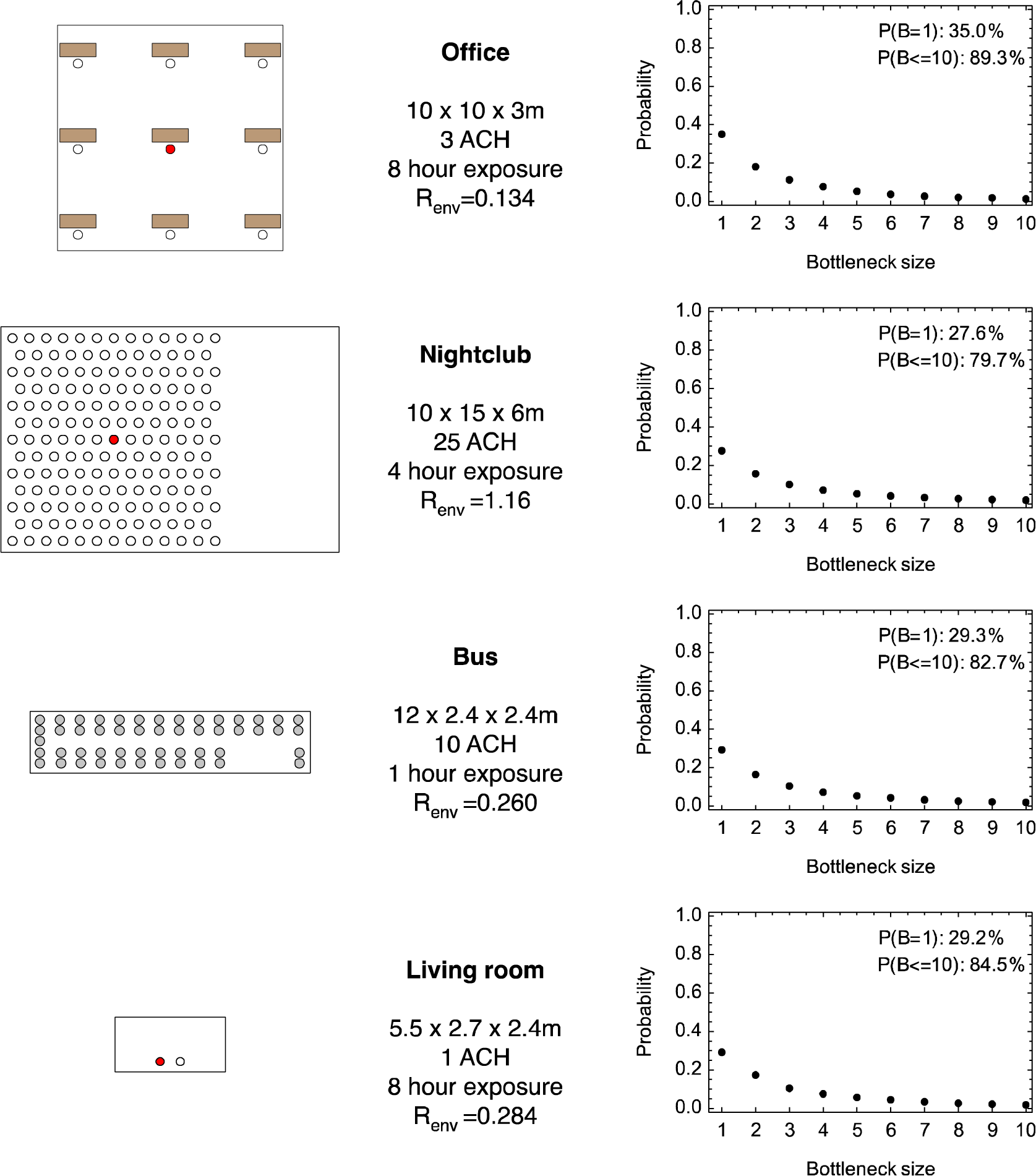
Bottleneck size distributions calculated for different scenarios. Maps show the layouts of different scenarios. A red dot indicates the location of an infected person, while a white dot indicates the location of an uninfected person. In the bus the location of the infected individual was chosen at random, with results being averaged across possible locations; gray dots show the locations of individuals. In our model individuals were assumed to remain stationary. Furniture did not affect the model and is shown for purely illustrative reasons. Data show the room dimensions. Ventilation levels are described by the number of air changes per hour (ACH). The value R_env_ describes the expected number of people infected in an environment during the modelled time of exposure.

Different environments in our model differed in the number of cases of infection we expected to occur. We denote by R_env_ the expected number of cases in an environment during the specific period of time for which that environment was modelled; this mirrors the epidemiological parameter R_0_, which describes the total expected number of infections caused by an infected individual during the entire course of infection^28^. Under default parameters our model predicts that an expected 1.16 infections would occur in the nightclub over a four-hour period; fewer infections occurred in the other environments.

Estimates of R_env_ were derived from an estimate of R_0_ and an assumption that the office environment, as we describe it, is typical of the many possible environments in which transmission might occur. Taking R_0_ to mirror SARS-CoV-2 transmission and assuming the office to be a typical environment implies an expected number of cases in the office. This number of cases, and the volume exposures in our model, together imply a specific effective viral load.

The effective viral load calculated by our model under default parameters was 9.7 x 10^8^ viruses per ml. Under these parameters the inhalation of an infectious particle is a rare event, but one which often leads to infection: Inhaling a single particle with initial radius 10μm would lead on average to four viruses forming part of the transmission bottleneck. The need for a particle to be inhaled is the key barrier preventing transmission (Supplementary Figure S1). The small bottleneck sizes we infer reflect the fact that larger particles were rarely inhaled, being disproportionately removed from the air by sedimentation (Supplementary Figure S2).

Our basic result, that most transmission bottlenecks are small, was robust to changes in our model parameters. The assumption in our model that the office is a typical transmission environment is represented by a parameter, φ_office_, being set to equal one. Increasing or decreasing this parameter reflects a belief that the office environment has a higher or lower than average propensity to be the location for viral transmission events: Changing this parameter has a direct effect on the expected number of infections in the office (Figure 3A), and upon the effective viral load (Figure 3B).

**Figure 3:**
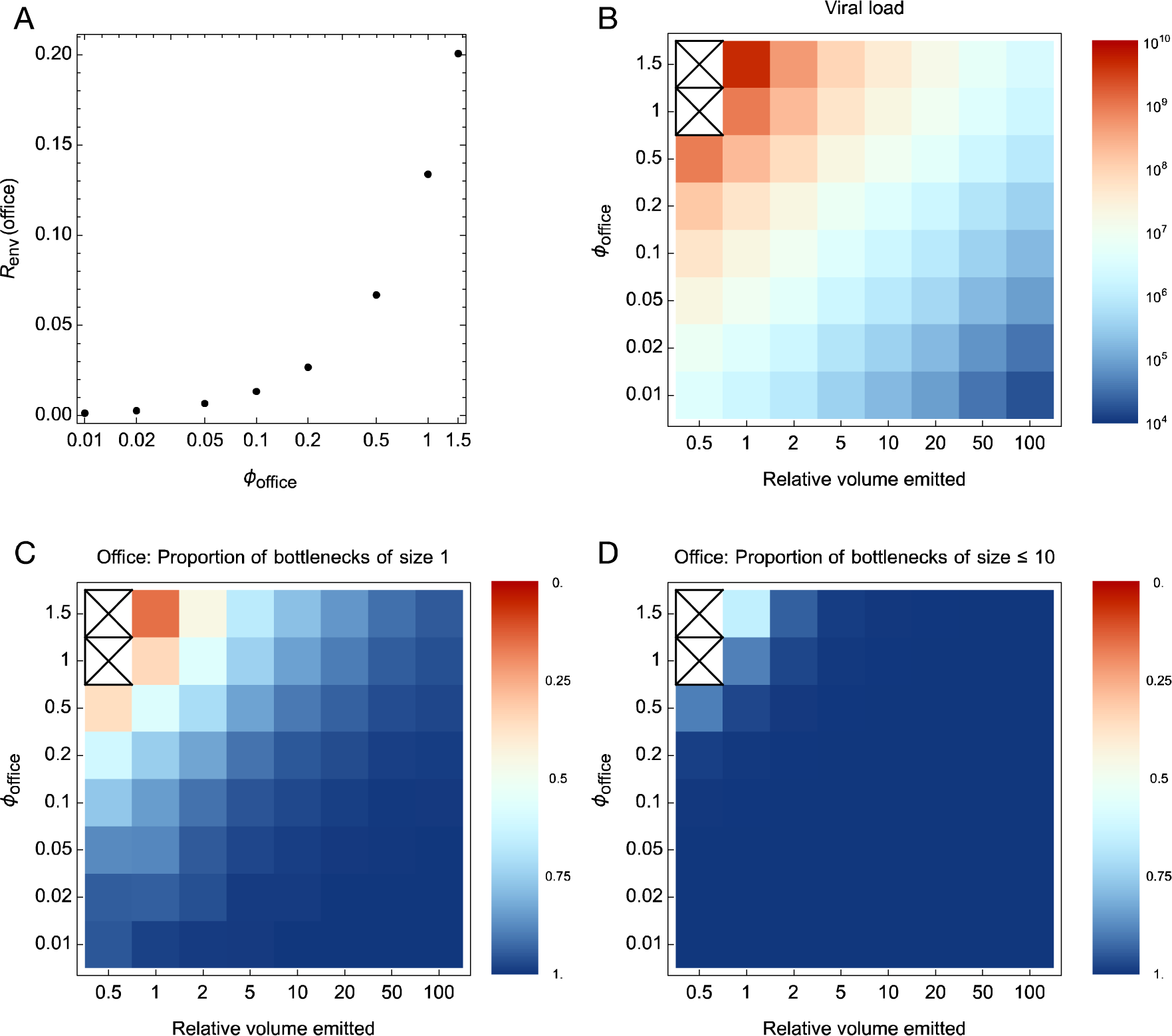
Sensitivity of our model to simple changes in model parameters. **A.** Changes in φ_office_, which describes the relative propensity of the office environment to be a location for viral transmission, are directly related to R_office_, which describes the expected number of infections taking place in the office during the time simulated in our model. **B.** The effective viral load, measured in copies per ml, increases with φ_office_, but decreases with higher emitted volume. An X indicates that the required number of infections could not be obtained for any effective viral load. **C.** Proportion of transmission bottlenecks involving a single viral particle. **D.** Proportion of transmission bottlenecks involving 10 viral particles or fewer.

The volume of particles emitted by the infected host is another parameter subject to uncertainty^29^. Where in our model an increased number of particles was emitted, a lower viral load was required to cause the same expected number of infections (Figure 3B). If we assume that a high volume of particles is emitted, our model produces a different paradigm for transmission. In that situation the inhalation of particles becomes a common event, although the decreased viral load means that inhaling a single particle rarely causes an infection (Supplementary Figure S3).

Although changes in our model parameters change some details of our output, the result that most transmission bottlenecks are caused by tight transmission bottlenecks was robust to changes in φ_office_ or in the total volume of particles emitted. Indeed, bottlenecks generated under our default parameters were larger than those obtained in most circumstances: Reducing φ_office_ to 0.1 led to 85% of transmission events involving a single virus. Among the combinations of parameters tested, the largest bottlenecks arose at a default level of particle emission and a value φ_office_ = 1.5: At these values only 16% of transmission bottlenecks in the office comprised a single virus, with 65% of comprising fewer than 10 viruses (Figure 3C,D). The extremely high effective viral load in this scenario, of 4.6 x 10^9^ viruses per ml, implies that infectious particles can carry large effective numbers of viruses, causing high population bottlenecks. However, as we discuss later, such a viral load is likely to be unrealistic; sets of parameters involving lower viral loads, and consequently tighter bottlenecks, are more likely. Patterns of bottlenecks derived for the office were replicated in other environments (Supplementary Figure S4). We note that some combinations of parameters are not self-consistent: low levels of particle emission cannot produce the highest numbers of cases of infection.

To this point we have shown that, across a range of scenarios in which the expected number of infections was constrained to a realistic value, transmission was predicted to generate tight transmission bottlenecks. In an attempt to generate larger transmission bottlenecks, we re-parameterised our model to simulate superspreading, whereby a ten-fold increase in the number of particles emitted was accompanied by a very high effective viral load of 2 x 10^9^ per ml. This change increased the number of cases of transmission, with on average more than 13 people infected in the nightclub (Figure 4). However, in three of the four environments, the most common transmission bottleneck size remained as one, with two thirds or more of transmission events still involving 10 or fewer viruses establishing infection. The one exception to this was the living room in which a broad distribution of bottleneck sizes was observed, peaking at a bottleneck of 8 viruses. The distinct result obtained for the living room reflects a greater combined exposure to viral particles at a unit distance (Supplementary Figure S5): Under superspreading conditions, a long period of time close to an infected person in conditions of poor ventilation led to a higher transmission bottleneck. We note that, in a population where superspreading occurs, it happens in a context where most infected individuals have lower infectivity. An extension of our model which incorporated a range infectivity values across a population led to the most common bottleneck size in the living room being one, even where the mean parameters matched our superspreading conditions (Supplementary Text, Supplementary Figures S10, S11).

**Figure 4:**
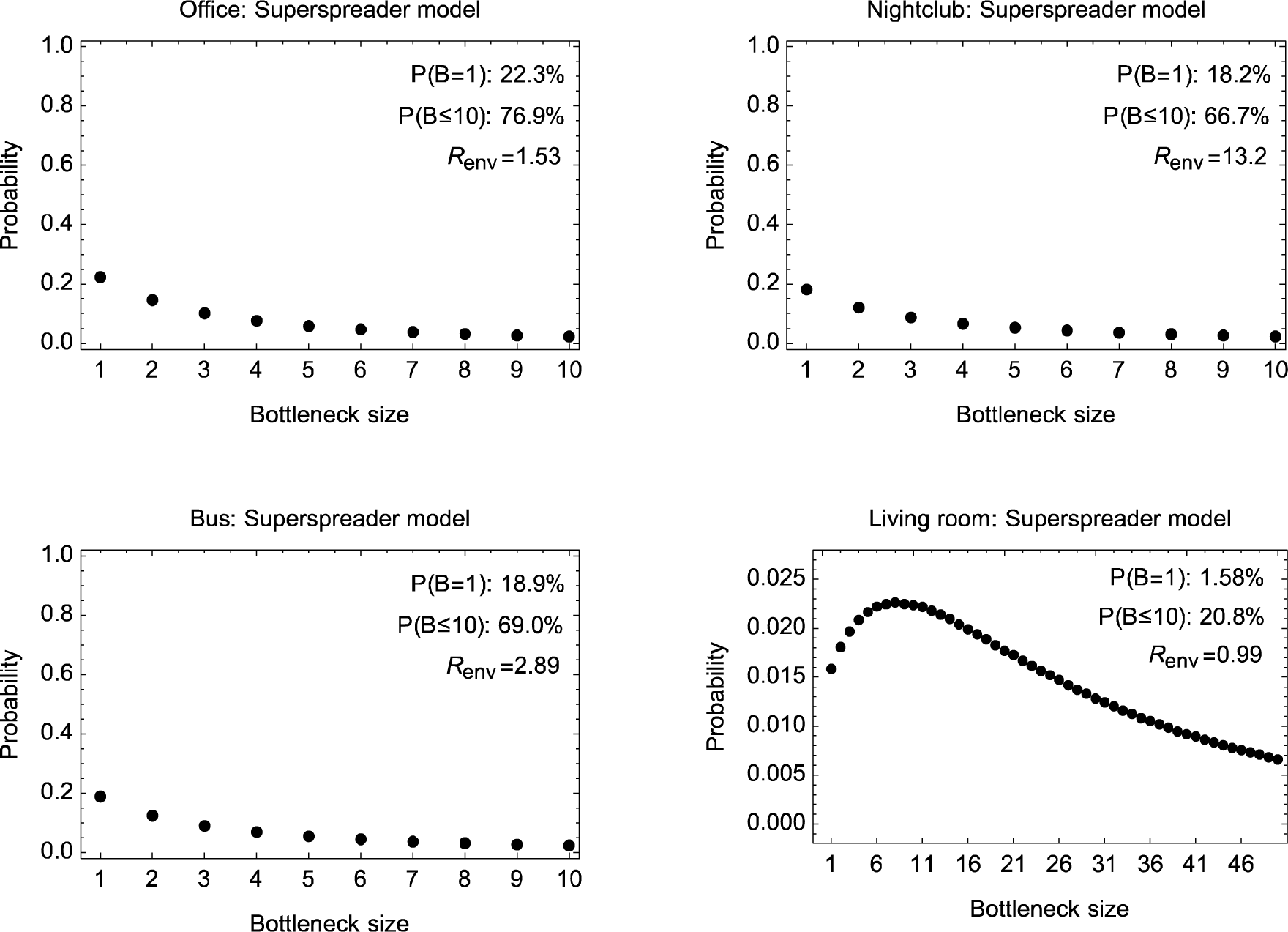
Transmission bottlenecks inferred under a superspreading model. The bottleneck distributions shown reflect an effective viral load of 2 × 10^9^ per ml and an extent of virus emissions 10-fold higher than our default model.

Our result that the majority of transmission events involve small numbers of viruses was replicated when we explored forms of particle emission other than coughing. Speaking and sneezing produce distributions of particles sizes distinct from that of coughing. In the case of speaking, the distribution, as measured via experiment, is not greatly dissimilar to that of coughing. While speaking may produce a different total volume of particles to coughing, the distribution of the sizes of those particles was not sufficiently different for our model to provide distinct results (Supplementary Figure S6). Relative to coughing, sneezing produces a much greater overall volume of particles than coughing, and has a distinct distribution skewed towards particles of larger sizes. Constraining a model of sneezing to produce the same expected number of infections for the office produced a distribution of transmission bottlenecks that was sharply skewed towards small bottleneck sizes (Supplementary Figure S7). The pattern of infection from the sneeze model is comparable to an extreme version of the coughing model in which the volume emitted is greatly increased: Very many particles are inhaled but a low viral load means that each particle contributes a small number of viruses to the bottleneck.

In a model describing the physical process of respiratory virus transmission, in which the expected number of people infected is constrained by reasonable parameters, tight transmission bottlenecks prevail. This result holds across a broad range of model parameters: We conclude that, wherever this process of virus transmission occurs, tight transmission bottlenecks are to be expected.

## Discussion

We have here used a modelling approach to describe the physical process of airborne virus transmission. Combining a model of particle emission and diffusion with a realistic rate of infection we infer that the physical process of transmission conspires to generate transmission events that involve tight bottlenecks, with 10 or fewer viruses founding infection. The exact distribution of bottleneck sizes inferred by our model depended upon the prevailing environment and the time spent within that environment. High transmission bottlenecks were observed given an extended period of close proximity to a person shedding a high volume of material with a very high viral load, and in a poorly ventilated environment. They also occurred in rare circumstances under our default model conditions, via exposure to a large droplet containing multiple viruses. However, as a rule, tight transmission bottlenecks were in the majority.

Our inferred distributions of bottleneck sizes reflect estimates from previous studies derived from viral genome sequence data. For example, a study of influenza virus suggested that between 28 and 31 (73 to 82%) of a set of 38 transmission events were likely to have been founded by a single virus^2,30^. The same study identified one case of infection which apparently involved a very high transmission bottleneck. Our work provides a physical rationale for the inference of tight transmission bottlenecks from previous studies that have used genome sequence data. However, in so far as our approach does not rely upon genomic data, our result is more generalisable, spanning for example cases of respiratory viral transmission for which genomic data have not yet been collected. In this sense our work is predictive: If there were to be an outbreak of a novel virus spreading by airborne transmission, our work predicts that transmission events involving that virus would involve small numbers of viral particles. Our model makes further predictions about the dependence of transmission bottlenecks upon the local environment: Where people spend extended periods of time close together in a poorly ventilated environment, such as in a domestic setting, transmission bottlenecks are likely to be statistically higher than in well-ventilated public settings.

In constructing our model we have made multiple simplifications. For example, we neglect effects arising from convection currents caused by individuals in a room^31^. Such effects have the potential to generate non-monotonic levels of exposure with distance from an infected person, as particles are carried up and across the ceiling before falling to a height at which they can be breathed in. Our model neglects effects arising from changes in humidity within a room^32^, and further neglects detailed description of ventilation, such as the placement of windows, ceiling vents, and air conditioning units. Although the generalised picture described by our model allows the consideration of broad classes of environments, such factors could potentially increase disparities in the exposures of individuals.

Another simplification in our model is the neglect of interactions between viruses, which could increase or decrease bottleneck sizes. Some interactions, such as those characterised by superinfection exclusion, are likely to reduce the number of cases of large transmission bottlenecks. In many cases of acute respiratory infection, a virus founding infection leads to the rapid growth of viral particles^33^, such that after a given amount of time, any subsequent infection will involve the addition of a tiny fraction of the current within-host population. This, alongside the triggering of innate host immune responses^34^, and other cellular interactions^35^, limits the window of time available for new viruses to infect a host. Other interactions between viruses, involving cooperation, have the potential to increase the proportion of bottlenecks involving multiple virions^36^. Where single virions contain incomplete functional genomes, more than one may be required for a cell to produce a complete genome^37,38^. A consideration of viruses with incomplete genomes would require a more nuanced definition of what is meant by a population bottleneck.

Our consideration of ranges of parameters reflects uncertainties within our model. For example, we have only a weak understanding of the extent to which the environment of our office simulation is representative of a typical environment in which transmission might occur. The effective viral load we infer under default parameters, of 9.7 x 10^8^ effective particles per ml, is very high even if every virus inhaled were to go on to found infection. Although we do not know what proportion of the viruses that are inhaled go on to found an infection, this value likely reflects too high an estimate of the extent of transmission in the office. In this sense our choice of default parameters is a conservative one, potentially over- estimating the size of bottlenecks at transmission.

In constructing our model we are aware of uncertainty in the literature describing distributions of particle sizes emitted via coughing, speaking and sneezing. While our method exploits experimental results, studies of these processes have historically used different methods and are not in perfect agreement. Again we note that under the very different circumstances representing coughing and sneezing in our model our basic result holds. While we would not be confident about combining these models to obtain a comprehensive and unified model of particle emission, our result that such distinct models produce similarly small bottleneck sizes supports the robustness of our conclusion.

Despite its limitations, the generalisability of our model and the reproducibility of our result across a broad range of scenarios provide what we feel is a compelling explanation for past observations of tight transmission bottlenecks in respiratory virus transmission. Where the number of cases of infection in a scenario are limited, as represented by a low value of R_env_, most people exposed to an infected person are not themselves infected, incurring an effective transmission bottleneck of zero. In a case where, following a cough or sneeze, infectious particles spread via diffusion, it is difficult to generate patterns of exposure which incorporate both cases of non-infection, and cases of infection that exclusively involve large bottlenecks. Particle diffusion, a process intrinsic to respiratory virus spread, leads to tight transmission bottlenecks.

## Methods

### Transmission mediated by the airborne spread of infectious particles

We built a model of the airborne spread of viruses within a room. Considering in the first instance SARS-CoV-2 infection, we modelled the behaviour of particles emitted by an infected individual with a cough, measuring the subsequent exposure of others in the room. Considering a single coughing event, we estimated the exposure E_e_(x,t) resulting from the cough at a distance x metres from the infected individual and a time t minutes after the cough, given that the individuals share an environment e.

In our model time was evaluated in units of minutes. We calculated the extent of exposure each minute after a cough, summing these exposures over time. We suppose that the infected individual coughs multiple times, at the times t_j_, and that the uninfected person i remains at distance x_i_ through time. Our model is evaluated on a discrete timescale of minutes, summing over time and discrete coughing events. The total exposure of person i after time T was calculated as

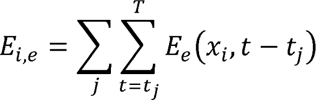

Our model assumes that the process of being exposed does not change the local level of exposure i.e. breathing in viruses does not significantly remove viruses from the air. We explore this assumption further in Supplementary Text.

### Diffusion model

To calculate exposure levels we consider the concentration c(r,x,t) of infectious material contained in particles of radius r at distance x from the infected person and time t. This concentration is altered by the emission of infectious particles into the environment, the spread of particles through space, the loss of particles via evacuation and sedimentation, and the degradation of viruses contained within particles. We write

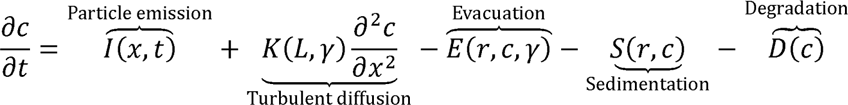

The diffusion of particles was modelled in a range of different environments. Environments differed in the number of uninfected individuals present, their relative locations, and the times for which they were exposed, and in the size of the environment and the extent to which that environment was ventilated.

### Emission of infectious particles

Coughing is a common symptom of SARS-CoV-2 infection^39^. Coughing leads to the emission of a distribution of particle sizes, which have been studied via a range of experimental means^19,40^. We modelled particles emitted from a cough as following a lognormal distribution, generating a distribution that covers particle sizes of diameters from zero upwards^20^. As such, we simulated particles with radii r ∈ {1, 2, …, 80} μm, with particles being emitted in quantities proportional to the function

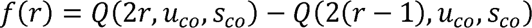

where Q is the cumulative distribution function of the lognormal distribution

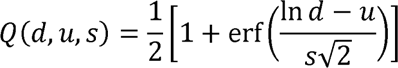

with the parameters u_co_ = 2.60269 and s = 0.693147^41^.

We assumed that the infected person coughed 10 times per hour at regular intervals^42^. Coughing emits particles with initially very high velocity, although this velocity falls rapidly within a fraction of a second^43^. We thus described a cough as creating an instantaneous cloud of Gaussian-distributed particles with zero velocity, located a mean distance of 20cm from the infected person, and with a standard deviation of 5cm. By default the volume of liquid emitted from a cough was set to equal 38 picolitres^44^.

For comparison we also investigated models of particle emission by coughing and sneezing. Following previous literature, speaking emitted a distribution of particle sizes given by the cumulative distribution function of the lognormal distribution Q(d,u_sp_,s_sp_), where u =2.77259 and s =0.597837^41^. We modelled speaking as creating an instantaneous Gaussian-distributed cloud of particles with zero velocity, at a mean distance of 10cm from the infected person, with a standard deviation of 2.5cm.

The sizes of particles emitted from a sneeze were described using a bimodal distribution described by from previous experimental work^21^. As such the proportion of particles of diameter d was set to be proportional to

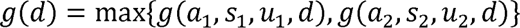

where

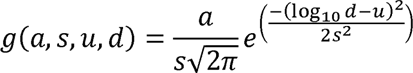

and the various parameters were given by a_1_=4.9359, s_1_=0.1479, u_1_=2.733$; a_2_=2.1649, s_1_=0.1483, u_1_=2.005. We modelled speaking as creating an instantaneous Gaussian- distributed cloud of particles with zero velocity, at a mean distance of 60cm from the infected person, with a standard deviation of 5.5cm. Distributions of the different particle sizes are shown in Supplementary Figure S8.

### Evaporation

Emitted particles evaporate over time, the removal of liquid leaving behind a smaller solid particle with radius approximately one quarter of that which was emitted^45–47^. This process occurs relatively quickly, with for example a droplet of size 20μm evaporating in under a second^47^, and a droplet of size 55μm evaporating within an estimated 14.5 seconds^48^. Considering a timescale of minutes, we assumed that the process of evaporation is short, such that a particle of radius r_0_ was instantaneously reduced to the new size r=r_0_/4.

### Turbulent diffusion

Once emitted, particles spread through the air via diffusion. Both Brownian motion and air turbulence potentially contribute to this, though at the size of particles we consider, it is likely that turbulent diffusion will dominate over Brownian motion^49^; our model considered only turbulent diffusion.

The extent of turbulent diffusion depends upon how well a room is ventilated, with more frequent replacement of the air in a room requiring a higher mean rate of particle movement. We used a model based upon the experimental measurement of air in a domestic environment^50^. We defined a characteristic length scale for a room by

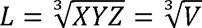

where V is the volume of the room in m^3^. Our model then links K, the turbulent diffusion coefficient, to L, and γ, the number of changes of the air in a room per hour:

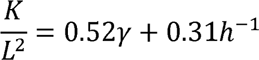

Within our model this becomes

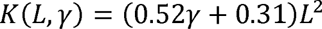

### Evacuation

We interpret the air change rate γ using the cutoff radius theory presented by Bazant et al^25^. Under this model, the evacuation rate is the same as the air replacement rate for droplets below a cutoff radius given by

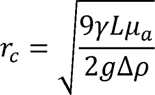

where μ_a_ is the dynamic viscosity of air, g is the acceleration due to gravity and Δρ is the difference in densities between water and air. Above the cutoff radius, the evacuation rate scales with 1/r^2^, with heavier particles being less subject to air movement. Where

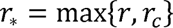

we have

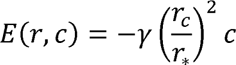

### Sedimentation

Emitted particles will be affected by gravity, with heavier particles falling to the floor more quickly than lighter particles according to Stokes’ Law. In our model we approximated this process by the simple removal of particles from the air over time. We followed a previous approach, which balanced diffusion and gravitational terms to approximate the time taken for a particle to fall to the ground^48^. We have

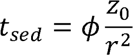

where z_0_ is the initial height of the particle, r is the particle radius, and φ is calculated as 0.85 x 10^-8^ ms. From this we derived the sedimentation term

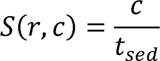

The initial height of particles z_0_ was defined according to whether individuals in an environment were standing or seated. We made the assumption that the floor absorbs particles, with no rebound being possible.

### Degradation of viruses

Viruses within emitted particles are intrinsically unstable, such that the number of infectious particles in a given droplet decays over time. An experimental study has suggested a half-life for SARS-CoV-2 of around 1.1 hours^23^. The viral titre in each droplet is therefore given by

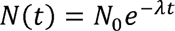

where N_0_ is the initial number of particles in the droplet and λ is the decay constant. To model a half-life of 1.1 hours, we set λ = 0.6301 h^-1^. We then have

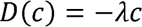

### Walls

We made the assumption that, upon hitting a wall, particles are absorbed, either impacting upon the wall due to electrostatic or inertial forces^51^ or being caught in downward convection currents leading to their deposition on the floor^31^.

### Units

We scaled the concentrations outputted from the diffusion model so that at time t=0 the total concentration c is equal to the volume in litres of a cough

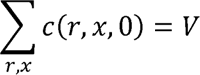

This implies scaling the c values so that for each r

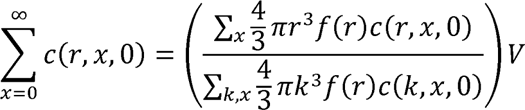

We set the volume V=38 picolitres, approximating to the extent of material contained in a cough^44^. The exposure at any given time was then specified by the equivalent volume of emitted material as it was at time t=0.

### Calculating individual exposures

The diffusion model described above was evaluated on a one-dimensional grid, calculating c(r,x,t) at positions on a linear grid x∈ X={0, 0.02, 0.04, …} centimetres from the infected person and at times t ∈ {0, 1, 2, …, 60} minutes following a cough. To calculate the exposure E_e_(x_i_,t) encountered by an individual we first approximate their location, finding the distance X_i_ as the value X_j_ ∈ X which minimises |x_i_-X_j_|. We next convert to polar coordinates and construct a box around the individual of width w metres, before considering exposure to particles of each different radius.

Considering a box of width w in polar coordinates, centred on a point at a distance X_i_ from the infected person, the total volume of infectious material it contains that is comprised of particles of original radius r was calculated as

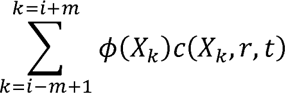

where

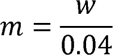

and

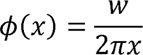

Is the fraction of the circle of radius x contained within the box. The volume of the box in metres cubed was then calculated as

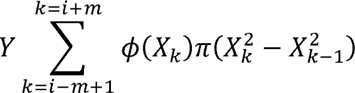

where Y is the height of the room in metres. Now, where A(t) is the volume of air breathed in by a person per minute, expressed in metres cubed, the exposure of that person at time t to particles of size r can be approximated by

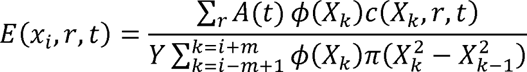

From this we calculate the total exposure of a person at x_i_ to particles of radius r:

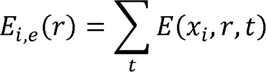

The expected number of particles to which that person is exposed may then be calculated as

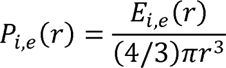

while the expected number of viruses that would cause infection that would be contained in a single particle of size r is in proportion to the particle volume

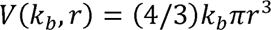

Where k_b_ is a parameter describing the effective viral load of particles at the point of emission; we calculate this parameter below.

Using the equations above we reproduced stochastic outcomes of transmission based upon exposures for each environment. Given an uninfected person, we calculated the number of particles of each radius to which they were exposed as a Poisson random variable with parameter P_i,e_(r). For each such particle we calculated the number of viruses forming the transmission bottleneck as a Poisson random variable with parameter V(r). For each scenario we repeated this process a minimum of 10^5^ times, generating a distribution of transmission bottleneck sizes for all cases of virus transmission (that is, where transmission involved at least one virus).

### Calculation of k_b_

Our model of particle emission and diffusion provides an expected level of exposure, measured in terms of a number of infectious particles, for people at different distances from the infected individual within the specified environment. In order to convert these exposures into transmission bottleneck sizes, we calculate an effective viral load k_b_. This parameter describes the number of viral particles per unit volume that would be expected to cause an infection, measured at the time of particle emission (i.e. prior to evaporation or degradation). This parameter is distinct from the physical viral load, as could be measured experimentally: Our parameter considers only the fraction of viruses that would overcome whatever barriers are in place in between being inhaled and founding a productive infection.

To calculate k_b_ we suppose that, of a total of n_e_ people exposed in the environment e, a mean of R people will be infected, where R ≤ n_e_. Infection occurs if and only if a person is infected by one or more viruses. Under our model a person is not infected either if they do not inhale any viral particles or if the particles they inhale do not contain any viruses founding infection. The inhalation of particles and the number of viruses contained by inhaled particles are both described by Poisson distributions. Therefore, the probability of not being infected by particles of radius r is given by

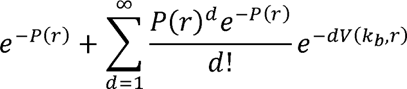

The expected number of infections in the environment e is therefore given by

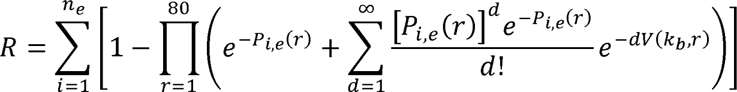

Considering the office environment, we used numerical optimisation to find a value of k_b_ in order to match a specific R value. In other environments R was calculated given knowledge of k_b_ and exposure values E_i,e_(r), which imply values of P_i,e_(r).

### Specification of R

In modelling infection, the statistic R_0_ the mean number of naive individuals who would be infected by an average infected individual^28^. This value is a composite term, averaged across a population of individuals who may have different levels of infectivity, and different numbers of contacts with other people in different environments. We decomposed this parameter, denoting by R_env_ the mean number of naive individuals infected by an average infected individual in a given environment during the time of our model. Setting R_env_ for any one environment fixes k_b_ and by consequence R_env_ for all other environments. Arbitrarily we chose the office environment, writing

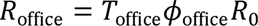

Here T_office_ is the expected fraction of time for which the infected person is in the office, as a fraction of the infectious period, and φ_office_ describes the propensity for infections to occur within the office environment as we have described it.

To calculate T_office_, we note that in our model, individuals spend eight hours in the same location, corresponding to half of the waking hours in that day. The fraction of the infectivity period was calculated in terms of a distribution of the times between symptom onset, published for SARS-CoV-2 early in the pandemic^52^, so that

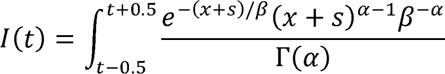

where α=8.124, β=1.572, and s=7.5. A 95% interval calculated for this distribution covers 18 days, with a mean of 5.35% of infections occurring on a single day. Using this value we derived a default value T_office_ = 0.5 x 0.0535 = 0.02675.

To comprehensively calculate φ_office_ would require a comprehensive understanding of all of the environments inhabited by people spreading infection. By default, we set φ_office_ = 1, assuming that our office model is perfectly typical of the heterogeneous environments people within a population inhabit. To model a virus similar to the original strain of SARS-CoV-2^53^ we set R_0_=5. In our results we explored values of φ between 0.01 and 1.5. Values of φ_office_ much greater than 1.5 were not accessible to our model given the other parameters; even in the limit of a high viral load R_office_ is limited by the requirement that an infected person be exposed to at least one infectious particle, and the default volume of liquid emitted by coughing. We note that relative to other workers during the pandemic, office workers did not have especially high rates of excess mortality^54^; a value of φ much greater than one seems unlikely.

Given a value for R_office_, we calculated k_B_ using the exposure values calculated for the office environment. This value of k_B_ was then applied to calculate bottleneck size distributions, and values R_env_, for other environments. As such, the effective viral load of emitted particles remains constant between environments.

### Environments

We modelled transmission within different environments, including an office, a bus, a nightclub, and a lounge. For each environment, our model was parameterised with the dimensions of the room in metres, X, Y, and Z, the number of people present, n_e_, and their locations, the air replacement rate γ, the length of time for which we assumed people were in the environment T, the volume of air breathed in per minute by an individual, A, and the height at which particles were emitted, z_0_. Parameters for each environment are shown in Table 1.

**Table 1:** Parameters defining each of the environments explored in our model. Parameters describe the dimensions of the room X, Y, and Z, the time spent in the given environment, T, the number of air changes per hour, γ, the number of uninfected people in the environment, n_e_, the volume of air breathed in per minute by uninfected individuals, A, here shown in units of litres, and the height at which particles were emitted, z_0_.

| Environment | X (m) | Y (m) | Z (m) | T (h) | $\gamma (h^{-1})$ | $n_e$ | A(L) | $z_0$ (m) |
| --- | --- | --- | --- | --- | --- | --- | --- | --- |
| Office | 10 | 3 | 10 | 8 | $3^{56}$ | 8 | $10^{55}$ | 1.1 |
| Nightclub | 10 | 6 | 15 | 4 | $25^{57}$ | 162 | $40^{55}$ | 1.6 |
| Bus | 2.4 | 2.4 | 12 | 1 | $10^{58}$ | 50 | $12.5^{59}$ | 1.1 |
| Living room | 5.5 | 2.7 | 3.7 | 8 | 1 | 1 | $6^{55}$ | 1.1 |

