## Supplementary Text and Figures for "The airborne transmission of viruses causes tight transmission bottlenecks"

1. *Removal of viruses from the air by inhalation*

Within our simulations we assume that the inhalation of viral particles does not affect particle concentration. That is, while the instantaneous exposure of an individual to infectious particles is affected by the local particle concentration, the presence of that individual does not affect the concentration. Further calculations showed that the extent to which viral particles would be removed from the air by inhalation was substantially less than the rate of removal via ventilation.

We assess the proportion of viral particles in a room removed per minute by breathing and by ventilation. As a simplifying assumption, we assume that the viral concentration is uniform throughout the room; our model did not incorporate a more complex reckoning for ventilation.

As such, we modelled the proportion of viral particles removed per minute by breathing as

$$A=\frac{n_{e}b_{e}}{v}$$

where n_e_ is the number of people in the room, b_e_ is the number of litres of air inhaled by each person, and v is the volume of the room in litres. For the living room we modelled b_e_ equal to 6 litres per minute, equal to a nominal resting rate, while in the nightclub we modelled b_e_ equal to 40 litres per minute, corresponding to moderate activity^1^. Other environments had intermediate values of this statistic.

In a similar way we modelled the proportion of viral particles removed per minute by ventilation as

$$p_{v}=1-{0.5}^{\gamma/60}$$

where γ is the number of air changes per hour. Here we model ventilation in a simple manner; assuming that an air change is equivalent to feeding in an amount of air equivalent to the volume of the room, thereby reducing the number of viral particles by one half.

Calculating these statistics, we found that in each environment ventilation was at least an order of magnitude more efficient at removing viruses than was breathing (Supplementary Figure S9). Our neglect of the latter term is therefore of the order of a correction to our ventilation rate.

2. *Extending our model to consider a distribution of infectivity values*

By default, our model considers transmission as involving a single individual, with the extent to which that person transmits the virus being modelled according to an R value. In a population, the extent to which a person is infectious will depend upon the time relative to the point of infection, and the individual characteristics of that person: studies have shown that not all individuals infected with SARS-CoV-2 are alike^2,3^. To model variation in the population we added a pre-factor to the exposure values Ei,e. This pre-factor was modelled as following a gamma distribution with parameters α=a and β=a^-1^, capturing a variety of potential distributions of host infectivity. We write

$$f\left( I,a \right)=\frac{I^{-a-1}a^{a}e^{-aI}}{\Gamma\left( a \right)}$$

In order to simulate a population of infected individuals we generated the centiles of the distribution. Where

$$F\left( I,a \right)=\frac{\Gamma\left( a,0 \right)-\Gamma\left( a,aI \right)}{\Gamma\left( a \right)}$$

We found values I_j_ for which F(I_j_,a) ∈ {0.01, 0.02, …, 0.99}. This gave the modified values

$$E_{i,j,e}\left( r \right)=I_{j}E_{i,e}\left( r \right)$$

The parameters E_i,j,e_ were normalised to have mean one, and used in place of E_i,e_ in our model; average bottleneck sizes were calculated over the parameter j. By default we set a=1, considering values of a ∈ {0.1, 0.5, 1, 2, 10} to explore the extent to which our results were sensitive to population heterogeneity. Results for a model calculated for different infectivity distributions under default parameters are shown in Supplementary Figure S10. Results calculated under superspreading conditions are shown in Supplementary Figure S11. We note that, while under a constant level of infectivity, superspreading conditions lead to the most common bottleneck size in the living room being equal to eight, where a distribution of infectivity values is considered, a bottleneck size of one is the most common. This result can be explained by the proportion of individuals in the modelled population who have low transmissibility: among infections caused by such individuals the most common bottleneck size is one.

**Supplementary Figures**

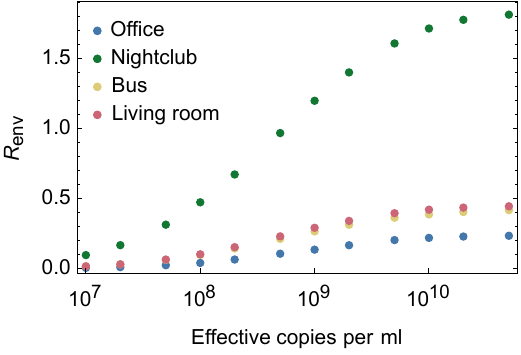

**Supplementary Figure S1: Sensitivity of our model to changes in the effective viral concentration.** In this figure other model parameters are kept at their default values. As the number of effective viruses per ml increases, the mean number of people infected R_env_ initially increases, but then reaches a plateau. Even where the number of viruses in a particle is extremely high, R_env_ is limited by the probability that people in each environment are not exposed to any infectious droplets. In total our model includes eight uninfected people in the office, 162 in the nightclub, 50 in the bus, and one in the living room.

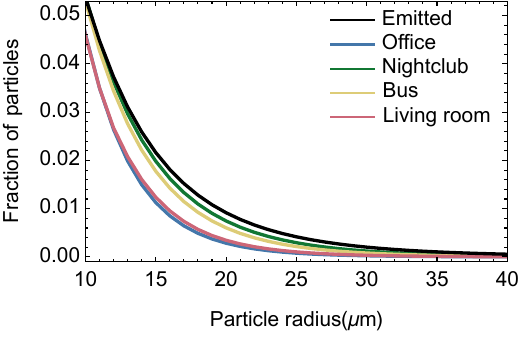

**Supplementary Figure S2: Distribution of the radii of particles emitted and inhaled.** Data are shown for particles at least 10um in size. The mean distribution is shown for each environment. In the office and living room environments lower rates of ventilation meant that there was more time for small particles to remain suspended in the air while larger particles were removed via sedimentation. This produces a greater bias towards smaller particles being inhaled.

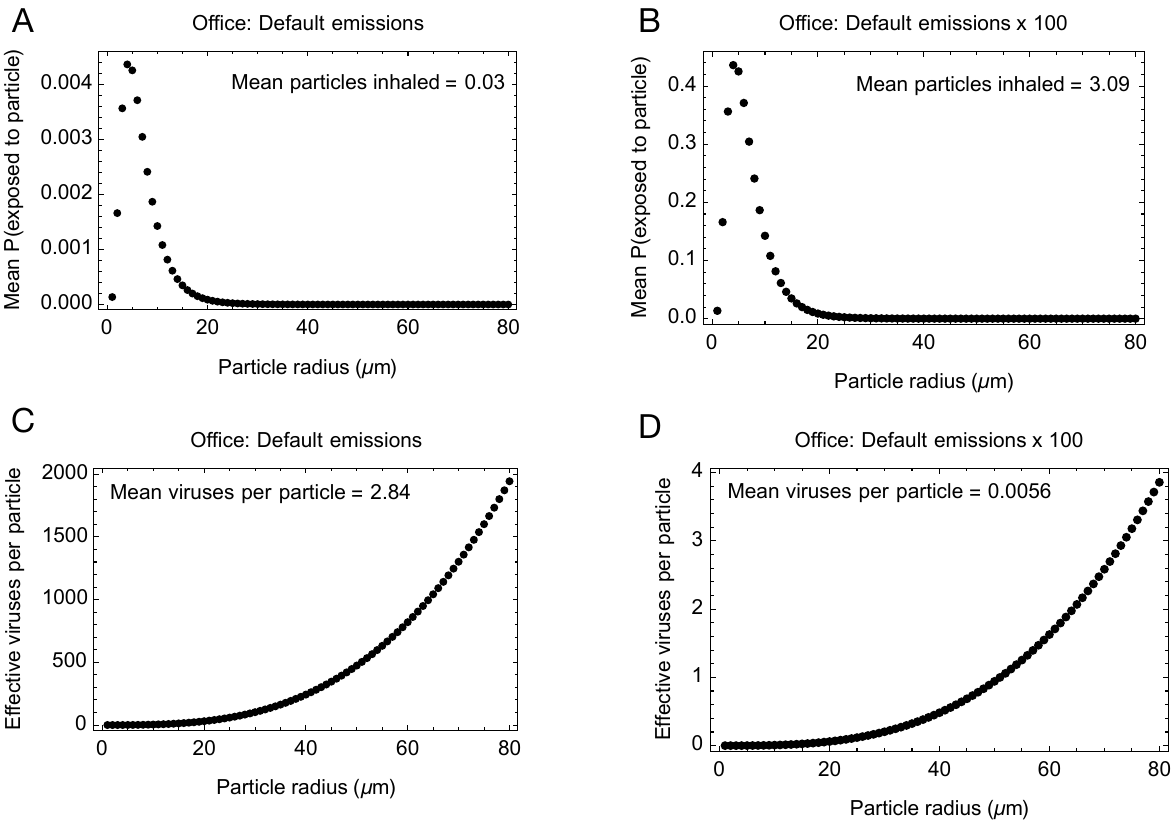

**Supplementary Figure S3: Effects of a change in the volume of emissions upon the model**. **A.** Mean probabilities of inhaling particles of different size calculated for people in the office environment under our default model parameters. **B.** Mean probabilities of inhaling particles of different size calculated for people in the office environment in a scenario where the number of emitted particles is increased 100-fold. **C.** Mean number of effective viruses per particle, plotted by particle radius, calculated for people in the office environment under our default model parameters. The calculated mean number of viruses per particle is calculated in a manner that accounts for the distribution of particle exposures. **D.** Mean number of effective viruses per particle, plotted by particle radius, calculated for people in the office environment in a scenario where the number of emitted particles is increased 100-fold.

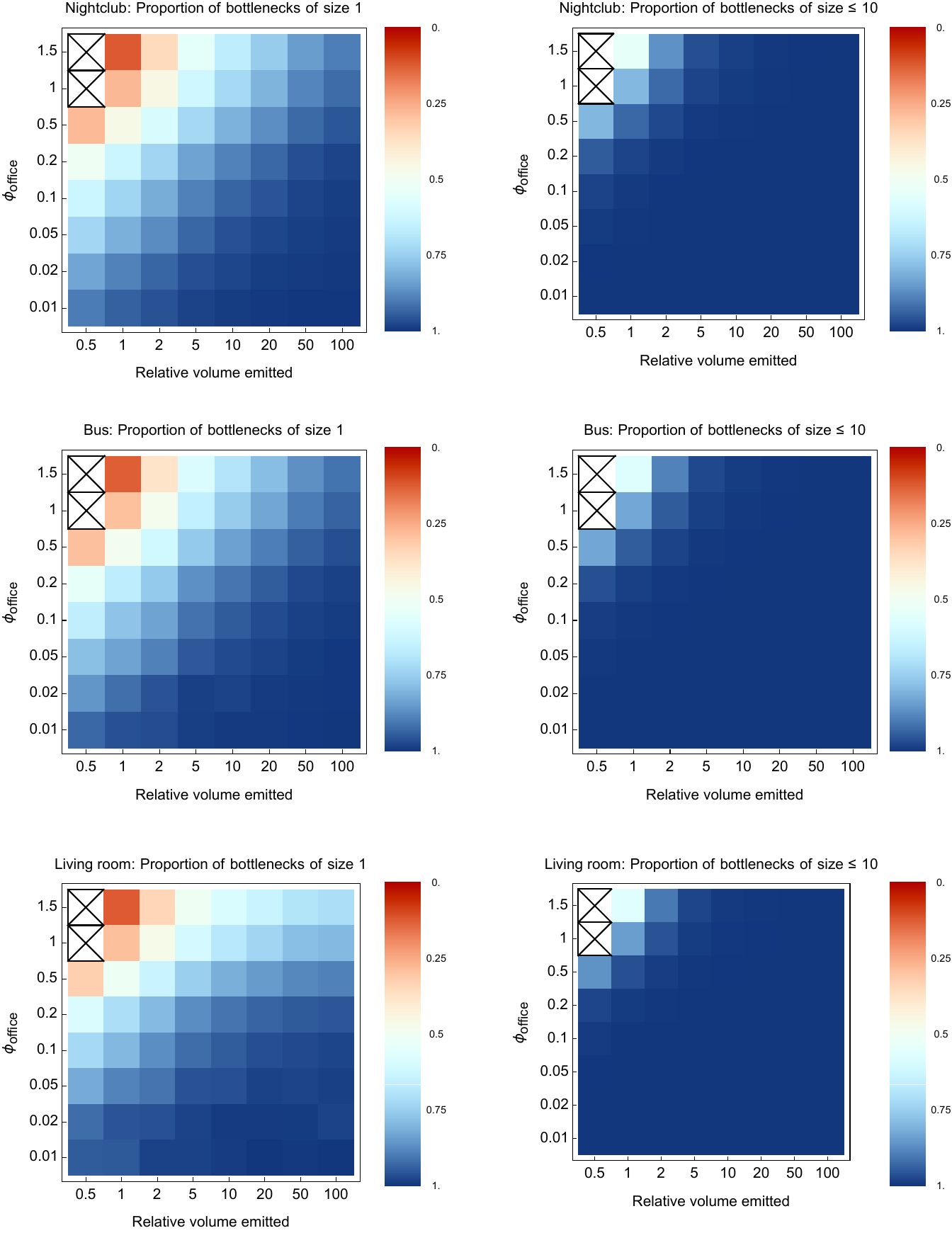

**Supplementary Figure S4: Bottleneck sizes for the nightclub, bus, and living room environments given basic changes in model parameters**. An X indicates that no result could be obtained for the relevant combination of parameters.

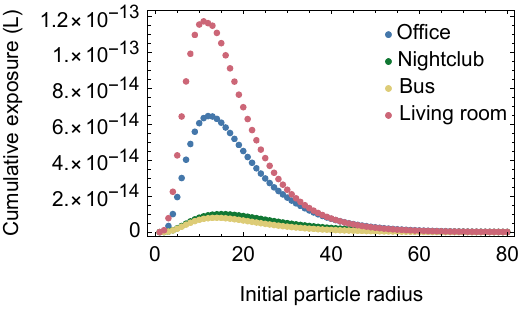

**Supplementary Figure S5: Combined exposure by particle size in each environment for a person at one metre distance from the infected individual**. Differences in exposure correspond to differences in ventilation in each environment, and in the length of time spent within an environment.

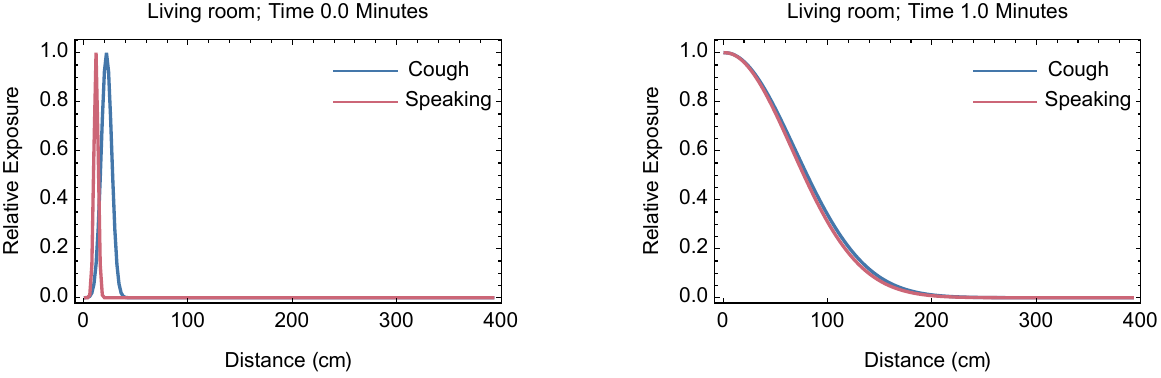

**Figure S6: Relative exposure to viral particles immediately following a cough or speaking event, and one minute after the event.** Distributions are shown for a cough and for speaking. After one minute the distributions are close to being indistinguishable.

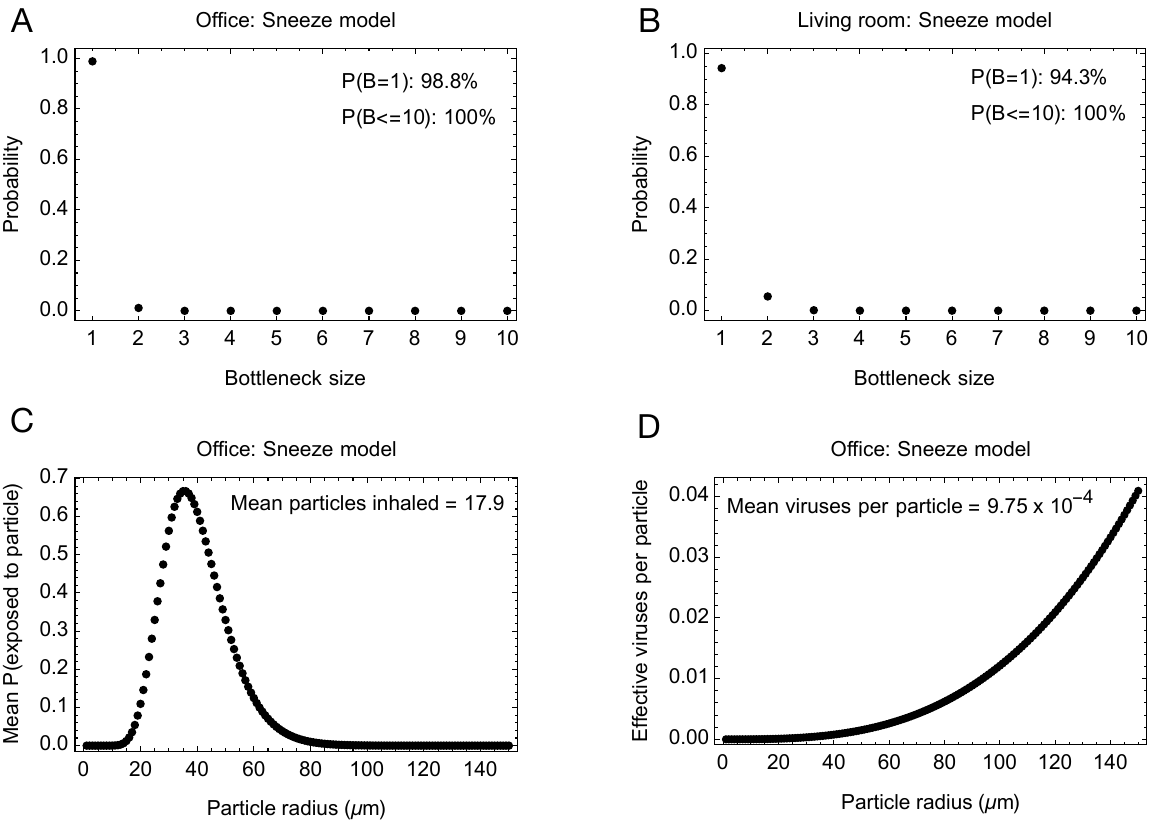

**Figure S7: Outputs from a model of sneezing.** Results were calculated for the office and living room environments, under the assumption that the infected person sneezed five times per hour. **A.** Transmission bottleneck distribution for the office environment. **B.** Transmission bottleneck distribution for the living room environment. **C.** Mean probabilities of inhaling particles of different size calculated for people in the office environment. Particles inhaled are larger than in the model of coughing. **D.** Mean number of effective viruses per particle, plotted by particle radius, calculated for people in the office environment. The calculated mean number of viruses per particle is calculated in a manner that accounts for the distribution of particle exposures.

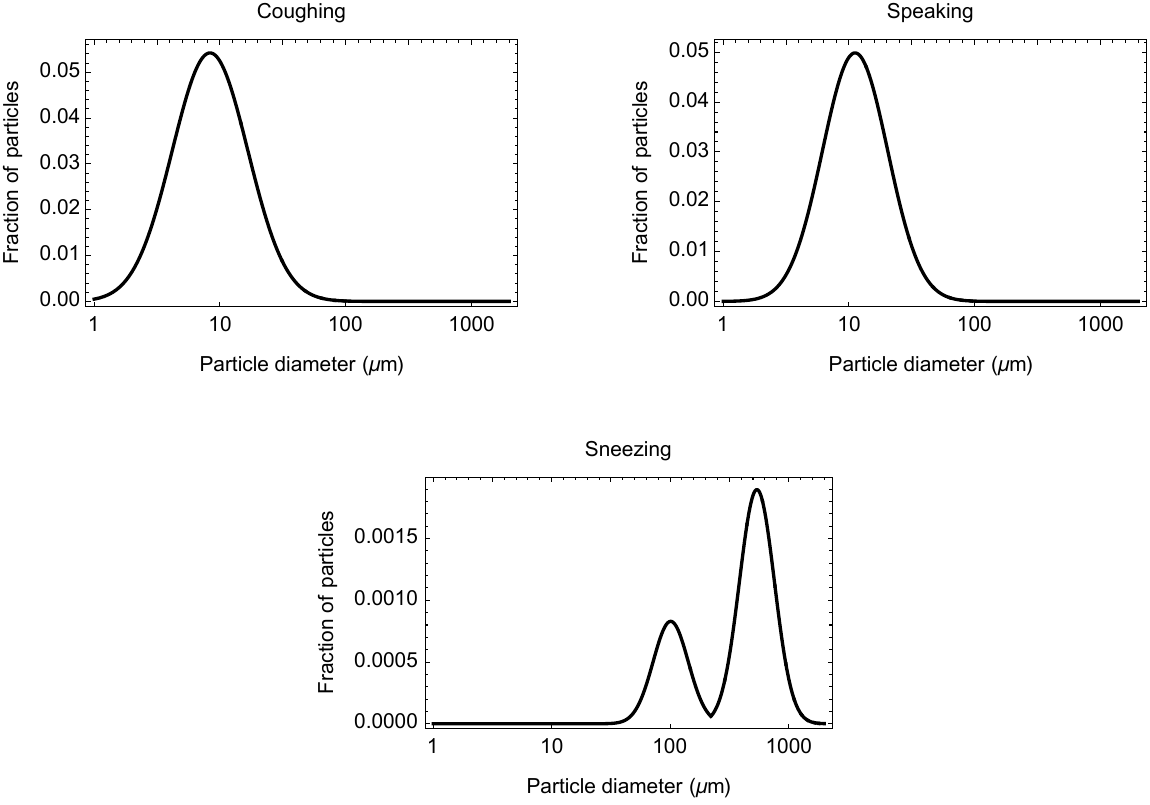

**Supplementary Figure S8: Experimentally-derived distributions of particles emitted via coughing, speaking, and sneezing, used in our model.**

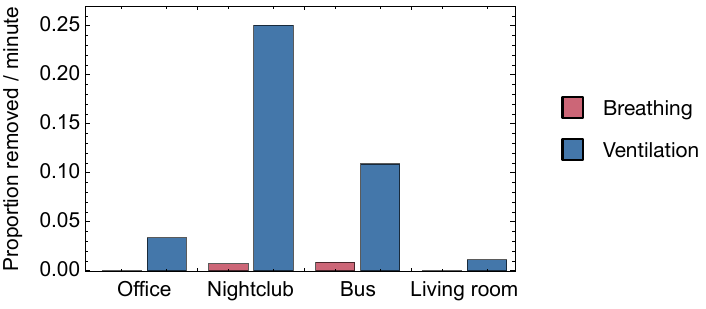

**Supplementary Figure S9: Proportion of viruses removed from the environment by breathing and by ventilation.** Values are shown as the proportion removed per minute.

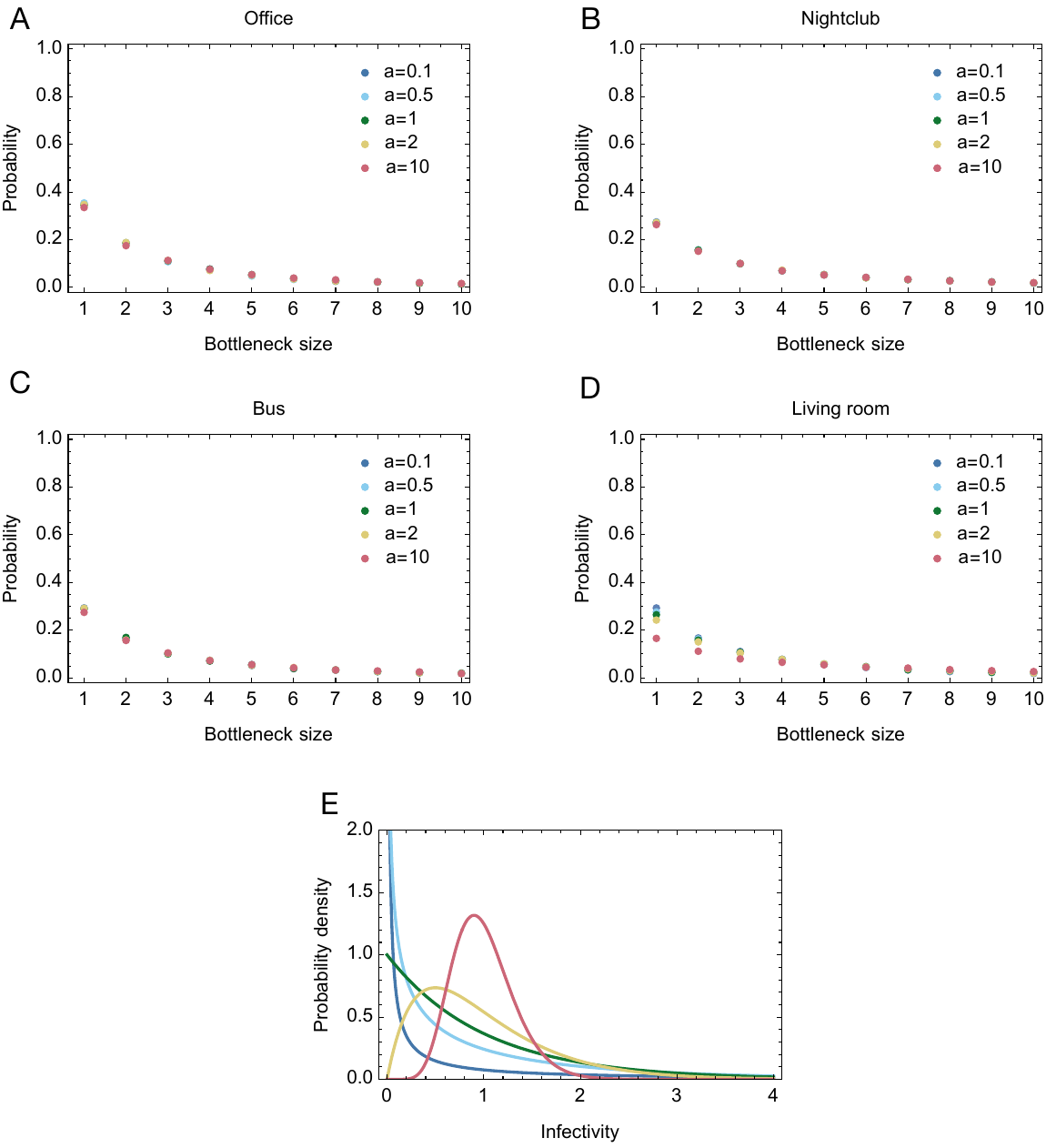

**Supplementary Figure S10: Distributions of bottleneck sizes derived under a distribution of infectivity values.** Other parameters were kept at their default values. Changes in the infectivity distribution did not lead to large changes in the inferred bottleneck sizes, albeit a slight increase in the number of cases with a bottleneck of 1 was observed as the extent of superspreading increased. Figure E shows the raw infectivity distributions.

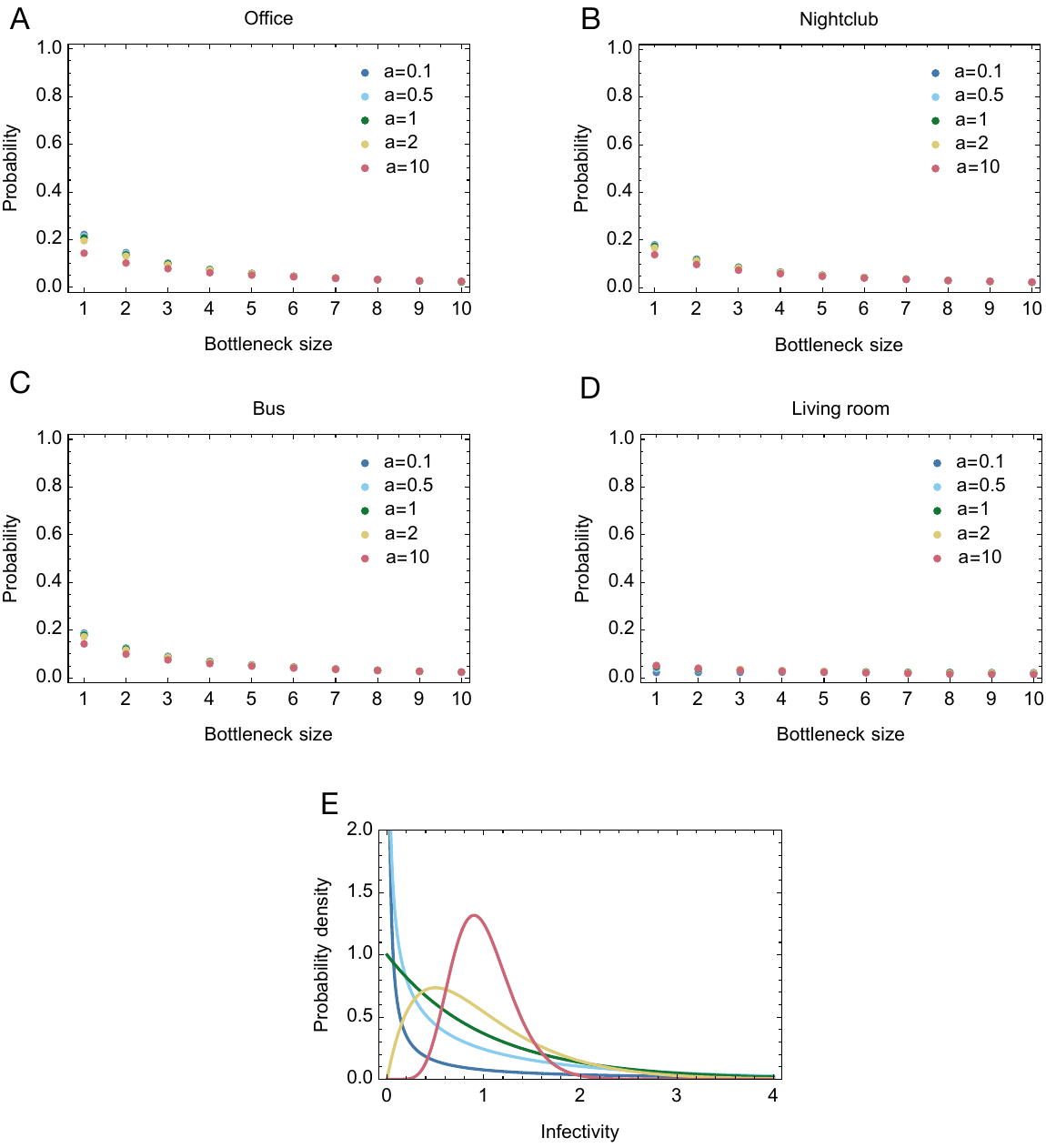

**Supplementary Figure S11: Distributions of bottleneck sizes derived under a distribution of infectivity values and under superspreading conditions.** An emitted volume 10 times the default for coughing was used, alongside an effective viral load of 2 x 10^9^ ml^-1^. Figure E shows the raw infectivity distributions.
